# Genomic diagnosis for children with intellectual disability and/or developmental delay

**DOI:** 10.1101/084251

**Authors:** Kevin M. Bowling, Michelle L. Thompson, Michelle D. Amaral, Candice R. Finnila, Susan M. Hiatt, Krysta L. Engel, J. Nicholas Cochran, Kyle B. Brothers, Kelly M. East, David E. Gray, Whitley V. Kelley, Neil E. Lamb, Edward J. Lose, Carla A. Rich, Shirley Simmons, Jana S. Whittle, Benjamin T. Weaver, Amy S. Nesmith, Richard M. Myers, Gregory S. Barsh, E. Martina Bebin, Gregory M. Cooper

## Abstract

**Background:** Developmental disabilities have diverse genetic causes that must be identified to facilitate precise diagnoses. We describe genomic data from 371 affected individuals, 309 of which were sequenced as proband-parent trios.

**Methods:** Whole exome sequences (WES) were generated for 365 individuals (127 affected) and whole genome sequences (WGS) were generated for 612 individuals (244 affected).

**Results:** Pathogenic or likely pathogenic variants were found in 100 individuals (27%), with variants of uncertain significance in an additional 42 (11.3%). We found that a family history of neurological disease, especially the presence of an affected 1^st^ degree relative, reduces the pathogenic/likely pathogenic variant identification rate, reflecting both the disease relevance and ease of interpretation of *de novo* variants. We also found that improvements to genetic knowledge facilitated interpretation changes in many cases. Through systematic reanalyses we have thus far reclassified 15 variants, with 11.3% of families who initially were found to harbor a VUS, and 4.7% of families with a negative result, eventually found to harbor a pathogenic or likely pathogenic variant. To further such progress, the data described here are being shared through ClinVar, GeneMatcher, and dbGAP.

**Conclusion:** Our data strongly support the value of large-scale sequencing, especially WGS within proband-parent trios, as both an effective first-choice diagnostic tool and means to advance clinical and research progress related to pediatric neurological disease.

## BACKGROUND

Developmental delay, intellectual disability, and related phenotypes (DD/ID) affect 1-2% of children and pose medical, financial, and psychological challenges [1]. While many are genetic in origin, a large fraction of cases are not diagnosed, with many families undergoing a “diagnostic odyssey” involving numerous ineffective tests over many years. A lack of diagnoses undermines counseling and medical management and slows research towards improving educational or therapeutic options.

Standard clinical genetic testing for DD/ID includes karyotype, microarray, Fragile X, single gene, gene panel, and/or mitochondrial DNA testing [2]. The first two tests examine an individual’s entire genome with low resolution, while the latter offer higher resolution but over a small fraction of a person’s genome. Whole exome or genome sequencing (WES or WGS) can provide both broad and high-resolution identification of genetic variants, and hold great promise as effective diagnostic assays [3].

As part of the Clinical Sequencing Exploratory Research (CSER) consortium [4], we have sequenced 371 individuals with one or more DD/ID-related phenotypes. 100 affected individuals (27%) were found to harbor a pathogenic or likely pathogenic (P/LP) variant, most of which were *de novo*. 16% of P/LP variants were identified upon re-analysis that took place after initial assessment and results return, supporting the value of systematic reanalysis of variant data to maximize clinical effectiveness. We also describe 21 variants of uncertain significance (VUS) in 19 genes not currently associated with disease but which are intriguing candidates. The genomic data we generated and shared through dbGAP [5], ClinVar [6], and GeneMatcher [7] may prove useful to other clinical genetics labs and researchers. Our experiences and data strongly support the value of large-scale sequencing for clinical and research progress related to pediatric neurological disease.

## METHODS

### IRB approval and monitoring

Review boards at Western (20130675) and the University of Alabama at Birmingham (X130201001) approved and monitored this study.

### Study participant population

Participants were enrolled at North Alabama Children’s Specialists in Huntsville, AL. A parent or legal guardian was required to give consent for all probands, and assent was obtained from those probands who were capable. Probands were required to be at least two years of age, weigh at least 9 kilos (19.8 lbs), and be affected with developmental and/or intellectual delays; more detailed information regarding enrollment, including phenotypic criteria, is provided in the Supplemental Methods.

### Whole exome and genome sequencing

Blood samples were sent for sequencing at the HudsonAlpha Genomic Services Laboratory (
http://gsl.hudsonalpha.org). Genomic DNA was isolated from peripheral blood and WES (Nimblegen v3) or WGS was conducted to a mean depth of 71X or 35X, respectively, with >80% of bases covered at 20X. WES was conducted on Illumina HiSeq 2000 or 2500 machines; WGS was done on Illumina HiSeq Xs. Reads were aligned and variants called according to standard protocols [8, 9]. A robust relationship inference algorithm (KING) was used to confirm familial relationships [10].

### WGS CNV calling

CNVs were called from WGS bam files using ERDS [11] and read Depth [12]. Overlapping calls with at least 90% reciprocity, less than 50% segmental duplications, and that were observed in five or fewer unaffected parents were retained and subsequently analyzed for potential disease relevance. All CNVs found within 5 kb of a known DD/ID gene, within 5 kb of an OMIM disease-associated gene [13], or intersecting one or more exons of any gene were subject to manual curation.

### Filtering and reanalysis

Using filters related to call quality, allele frequency, and impact predictions, we searched for rare, damaging *de novo* variation, or inherited X-linked, recessive, or compound heterozygous variation in affected probands, with modifications for probands with only one (duos) or neither (singletons) biological parent available for sequencing.

Potential secondary variants (i.e., medically relevant but not associated with the proband’s DD/ID) were also sought within parents. We assessed variants in 56 genes flagged by the American College of Medical Genetics and Genomics (ACMG) as potentially harboring medically actionable, highly penetrant genetic variation [14], those associated with recessive disease in OMIM [13], and carrier status for *CFTR, HBB,* and *HEXA*.

We also searched for all those variants listed as pathogenic or likely pathogenic in ClinVar [6], regardless of inheritance or affected status. Further details for variant annotation and filtration are supplied in Supplemental Methods.

For reanalysis, variants were reannotated with additional data, including updated versions of ClinVar [6], ExAC [15], DDG2P [16], and gene or variant lists identified in publications related to DD/ID genetics [17–19], and refiltered as described above and in Supplemental Methods. Candidate variants found in genes that were either not known to associate with disease or were found in individuals with phenotypes dissimilar from previously reported associations were submitted to GeneMatcher (
https://genematcher.org/) [7].

### Variant classification

Variants were classified into one of five categories: pathogenic, likely pathogenic, VUS, likely benign or benign. Our study began prior to publication of the formal classification system proposed by the ACMG [20], although our evidence and interpretation criteria are conceptually similar. Multiple lines of evidence, with mode of inheritance, allele frequency in population databases, and quality of previously reported disease associations weighing most heavily, are required to support assignments of pathogenicity. The Supplemental Methods contains a detailed description of our assertion criteria, and these criteria are also available via ClinVar [6]. The key annotations, including mode of inheritance, allele frequencies, PubMed identifiers, and computational inferences of variant effect, used to support the disease relevance of each variant are supplied in Table S2.

### Variant validation

WES and WGS were carried out under a research protocol and were not completed within a CAP/CLIA laboratory. All variants found to be medically relevant and returnable were validated by Sanger sequencing in an independent CLIA laboratory (Emory Genetics Laboratory) before being returned to participants, although these validated variant results are not CLIA-compliant as the input DNA was originally isolated in a research laboratory.

### Analysis of trios as singletons

For probands subjected to WGS as part of trios, we removed parental genotype information from their associated VCFs and subsequently filtered to identify variants that are expected to be extremely rare in the general population and/or affect genes known to associate with disease (Figure 2, Table S3). Scores from the Combined Annotation Dependent Depletion (CADD) algorithm [21] were subsequently used to rank P/LP variants within the filtered variant subset from each relevant proband. See Supplemental Methods for details.

**Figure 1:**
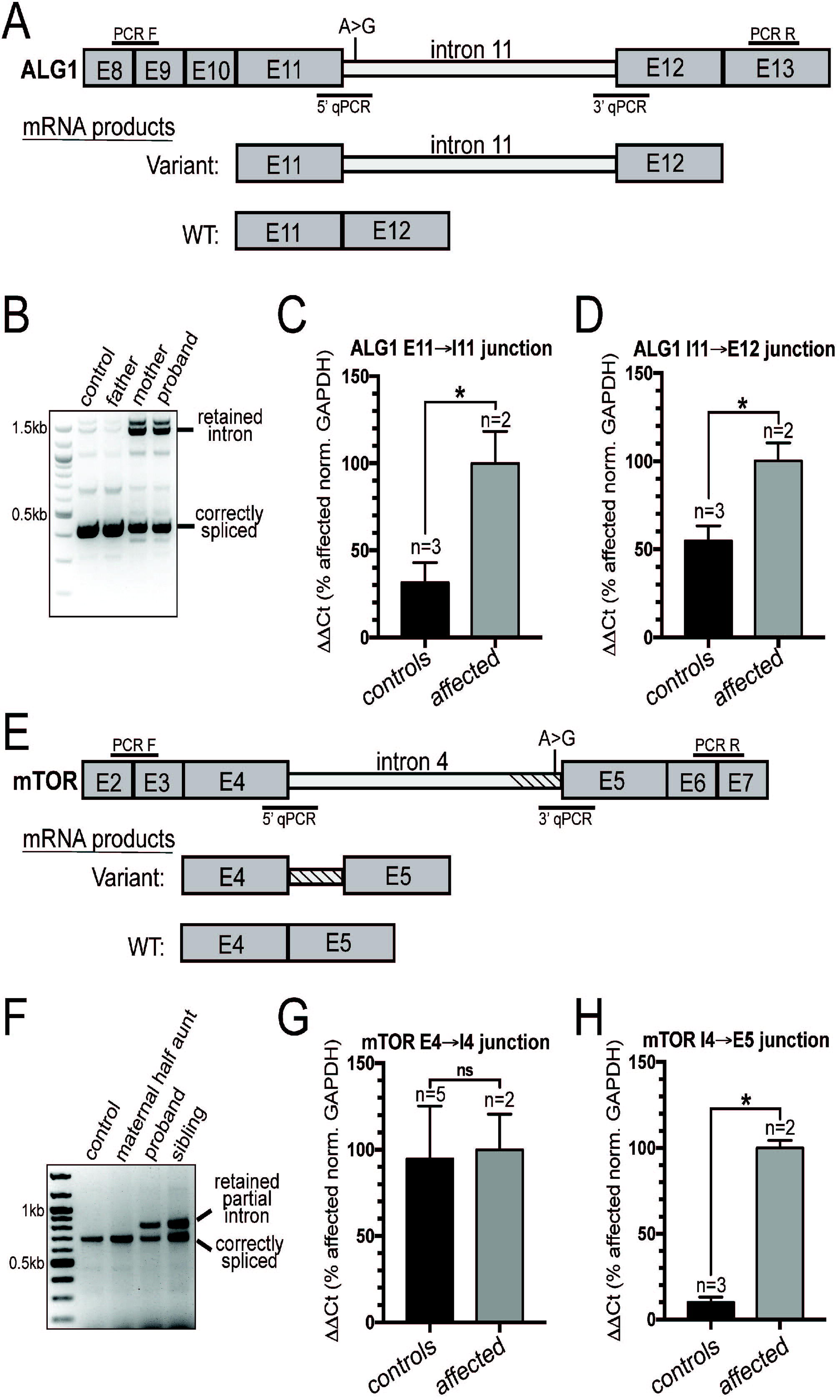
Intronic variants in *ALG1* and *MTOR* disrupt splicing and introduce early stop codons. (A) Diagram showing the region of *ALG1* surrounding the variant found in the proband and mother, an A>G transition 3 nucleotides downstream from the splicing donor site of intron 11. E=exon. (B) PCR F and PCR R indicate the position of the oligos used to amplify the region from patient derived cDNA. The control sample is from the RNA extracted from blood of an unrelated individual that did not harbor the variant. Control reactions lacking RT were also performed and did not show the PCR product containing the fully retained intron (data not shown). (C and D) qPCR analysis shows that the variant leads to inclusion of the entire intron 11. Controls are two unrelated individuals and the father of the proband. The affected individuals are the proband and mother. (E) Diagram showing the region of *MTOR* surrounding the variant, an A>G transition 2 nucleotides upstream of the splicing acceptor site. E=exon. (F) The region surrounding intron 4 was amplified using PCR F and PCR R (position indicated in E), and shows partial retention of the intron. The retained partial intron was not detected in control reactions lacking RT (data not shown). (G and H) qPCR from blood RNA shows that the 5’ splice site is not affected by the variant, but that the 3’ acceptor site is, leading to partial retention (134bp) of intron 4. Controls included unrelated individuals and the maternal half aunt of the proband. Affected individuals are the proband and half-sibling. For all qPCR analyses RNA was extracted from blood and ΔΔC_T_ values were calculated as a percent of affected individuals and normalized to *GAPDH*. The sequences of all oligos used are found in Table S7.

**Figure 2:**
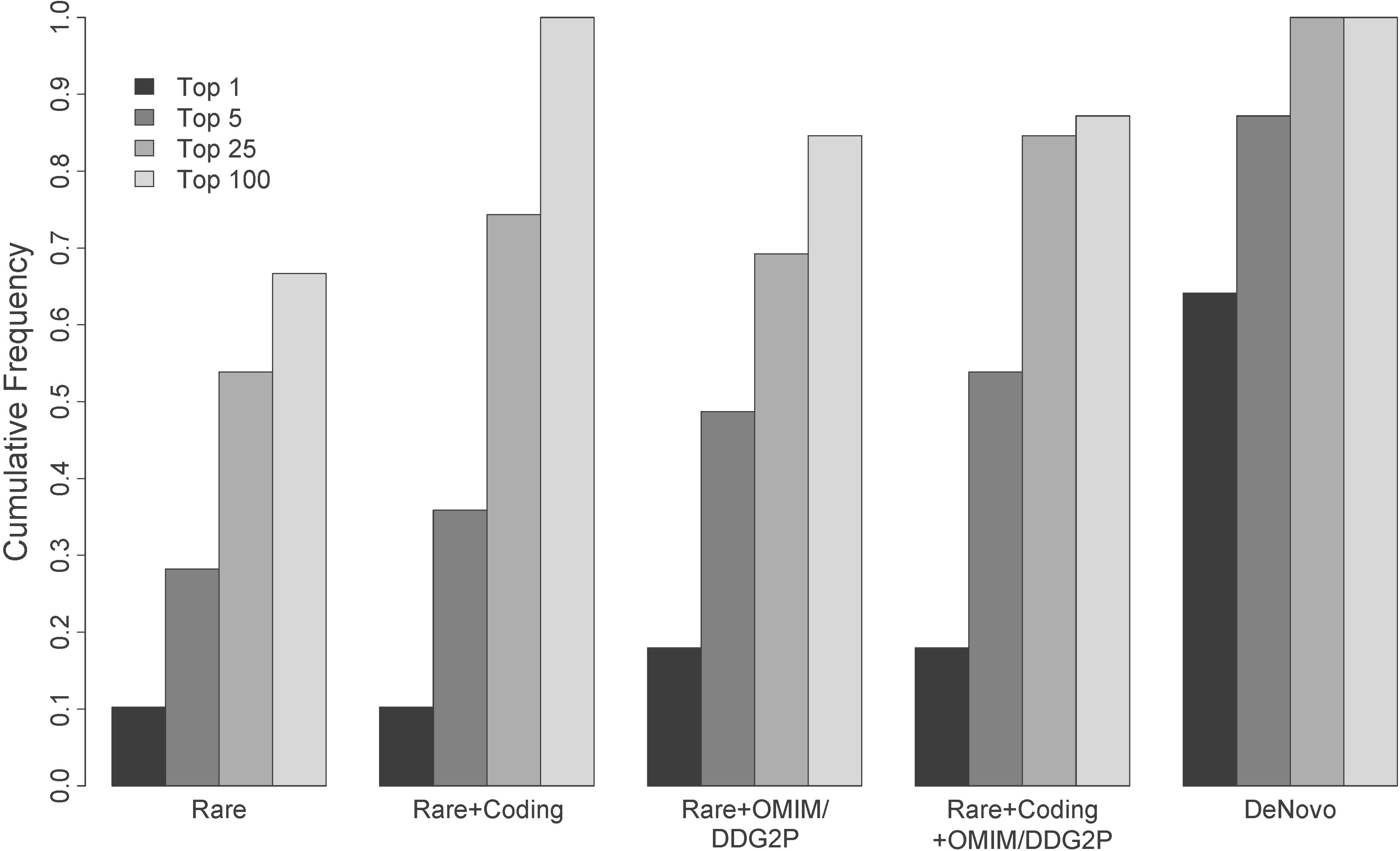
Ranks of pathogenic/likely pathogenic variants filtered without parental data relative to trio-defined *de novo* events. Most pathogenic/likely pathogenic variants, even under models that only consider population frequencies (e.g., “Rare”), rank (based on CADD) among the top 25 hits in a patient, and many rank as the top hit. Restrictions to rare coding variants and/or those affecting OMIM/DDG2P [13, 16] genes further enrich for causal variants among top candidates, making diagnosis feasible without parents.

### Functional Assays

RNA isolation, cDNA synthesis, qPCR and western blotting were conducted according to standard protocols. Details are provided in Supplemental Methods.

## RESULTS

### Demographics of study population

We enrolled 339 families (977 individuals total) with at least one proband with an unexplained diagnosis of a DD/ID-related phenotype (see “Study participant population” in Supplemental Methods). 284 participating families were enrolled with both biological parents. 261 of these families had one affected proband, while 21 families had two affected probands, and an additional two families had three affected probands. As each proband (including siblings within a family) was used to anchor a proband-parent “trio” as an analytical unit, our study includes a total of 309 trios from 284 families. We also enrolled 35 proband-parent “duos” that included one proband and one biological parent. Additionally, we enrolled two families with one biological parent and two affected probands (4 “duos”), and one duo family with three affected probands (3 “duos”), leading to a total of 42 “duos” from 38 families. Finally, we enrolled 17 “singleton” families in which no parents were available for testing; for 14 of these only one proband was tested, and in three families, two affected siblings were sequenced (total of 20 “singleton” probands).

During the course of this study, a decision to replace WES with WGS was made. In total, WES was performed on 365 individuals (127 affected) and WGS was performed on 612 individuals (244 affected). WES and WGS were sequenced to an average depth of 71X and 35X, respectively, with >80% of bases covered ≥ 20X in both experiment types. DNA from probands subjected to WES was also analyzed via a SNP array to detect copy-number variants (CNV) if clinical array testing had not been previously performed.

The study population had a mean age of 11 years and was 58% male. Affected individuals displayed symptoms described by 333 unique HPO [22] terms with over 90% of individuals displaying intellectual disability, 69% with speech delay, 45% with seizures, and 20% with microcephaly or macrocephaly. 18% of affected individuals had an abnormal brain magnetic resonance imaging (MRI) result and 81% of individuals had been subjected to genetic testing prior to enrollment in this study (Table 1).

**Table 1:** Pathogenic/Likely pathogenic rates by clinical annotation and family structure among the 371 DD/ID-affected individuals

| **CHARACTERISTIC** | **% Individuals**<br>**(No. of individuals)** | **% individuals with**<br>**P/LP result**<br>**(No. of individuals)** |
| --- | --- | --- |
| **AGE OF INDIVIDUAL** |  |  |
| 2-5 years | 25.8% (96) | 27.1% (26/96) |
| 6-12 years | 44.5% (165) | 25.4% (42/165) |
| 13-18 years | 16.5% (61) | 32.8% (20/61) |
| 19-40 years | 12.7% (47) | 32.0% (15/47) |
| >40 years | 0.54% (2) | 0.00% (0/2) |
| Average age of subject (age range) | 10.56 (2 to 54 years) |  |
| **SEX OF INDIVIDUAL** |  |  |
| Male | 57.7% (214) | 24.3% (52/214) |
| Female | 42.3% (157) | 32.5% (51/157) |
| **CLINICAL SPECIFICS** |  |  |
| Intellectual disability (mild, moderate, severe)<br>(HP:0001256, HP:0002342, HP:0010864) | 92.7% (344) | 28.5% (98/344) |
| Speech delay (HP:0000750) | 68.7% (255) | 27.1% (69/255) |
| Seizures (HP:0001250) | 45.3% (168) | 30.9% (52/168) |
| Facial dysmorphism (HP:0001999) | 30.2% (112) | 29.5% (33/112) |
| Autism spectrum disorder (HP:000729) | 25.6% (95) | 18.9% (18/95) |
| Hypotonia (HP:0001252) | 20.2% (75) | 34.6% (26/75) |
| Positive Brain MRI | 17.5% (65) | 28.1% (18/64) |
| Macrocephaly (HP:0000256) | 9.70% (36) | 25.0% (9/36) |
| Microcephaly (HP:0000252) | 9.16% (34) | 47.0% (16/34) |
| ADHD (HP:0007018) | 7.28% (27) | 25.9% (7/27) |
| Failure to thrive (HP:0001508) | 5.90% (22) | 27.3% (6/22) |
| Short stature (HP:0004322) | 4.85% (18) | 44.4% (8/18) |
| **PREVIOUS GENETIC TESTING** |  |  |
| Microarray | 59.8% (222) | 27.5% (61/222) |
| Single Gene/Gene Panel | 38.3% (142) | 30.3% (43/142) |
| Karyotype | 29.1% (108) | 36.1% (39/108) |
| Fragile-X | 27.2% (101) | 27.7% (28/101) |
| Mito DNA Screen | 7.55% (28) | 25.0% (7/28) |
| **FAMILY STRUCTURE** |  |  |
| Trio | 83.3% (309) | 29.1% (90/309) |
| Duo | 11.3% (42) | 19.0% (8/42) |
| Singleton | 5.4% (20) | 15.0% (3/20) |
| MRI, magnetic resonance imaging |  |  |

### DD/ID-associated genetic variation

WES and WGS data were processed with standard protocols to produce variant lists in each family that were subsequently annotated and filtered, and filtered variant lists were subject to manual review (see Methods). KING, a robust relationship inference algorithm, was used to confirm familial relationships [10]. Variant pathogenicity was classified based on allele frequency, inheritance status, published reports, computational deleteriousness predictions, and other sources of evidence; these assertion criteria are described in detail in the Supplemental Methods. All variants described here were confirmed by Sanger sequencing (see Methods) in probands and available family members before being returned to participants.

100 (27%) of the 371 probands had P/LP variants, while an additional 42 (11.3%) harbored a VUS (Table 2). Given that most probands had been previously tested via microarray prior to their enrollment in this study, large CNVs classified as a VUS or greater were detected in only 11 affected individuals (Table 2; Table S2; Figure S1).

**Table 2:** Results of exome and/or genome sequencing for 371 DD/ID-affected individuals

| **Assay**<br>**(Affected individuals)** | **SNV/indel** |  |  | **CNV** |  |  |
| --- | --- | --- | --- | --- | --- | --- |
|  | **Pathogenic** | **Likely pathogenic** | **VUS** | **Pathogenic** | **Likely pathogenic** | **VUS** |
| **Exome (127)** | 20.4% (26) | 9.4% (12) | 11.0% (14) | 1.6% (2)* | 0% (0) | 0% (0) |
| **Genome (244)** | 18.0% (44) | 4.1% (10) | 10.2% (25) | 2.0% (5) | 0.4% (1) | 1.2% (3) |
| **Exome and genome**<br>**(Total individuals: 371)** | 18.9% (70) | 5.9% (22) | 10.5% (39) | 1.9% (7) | 0.3% (1) | 0.8% (3) |

\* Identified by microarray

Most (76%) P/LP variation occurred *de novo,* while 12% of individuals inherited P/LP variants as compound heterozygotes or homozygotes (Figure S2A). An additional 5% were males with an X-linked maternally inherited P/LP variant. Finally, 7% of participants who harbored a P/LP result were sequenced with one or no biological parent and thus have unknown inheritance (Figure S2A). Most P/LP variants were missense mutations (52%), while 39% were nonsense or frameshift, 7% were predicted to disrupt splicing, and 2% led to inframe deletion (Figure S2B). Variants that were classified as a VUS or greater were identified in 97 genes, excluding large CNVs, with variants in 23 (24%) of these genes observed in two or more unrelated individuals (Tables S1 and S2).

### Pathogenic/likely pathogenic variant rates across families of varying structure and phenotypic complexity

Affected individuals were categorized into one of three analytical structures based on the number of parents that were sequenced along with the proband(s): proband-parent trios (309); duos with one parent (42); and proband-only singletons (20). A P/LP result was found in 29.1% of trio individuals, 19% of duo individuals, and 15% of singletons (Table 1).

We believe that at least some of the decline in P/LP variant yield in duos and singletons reflects the analytical benefits of trio sequencing to efficiently highlight *de novo* variation. However, given that one or both biological parents were unavailable or unwilling to participate in duo or singleton analyses, the P/LP rate comparisons among trios/duos/singletons may be confounded by other disease-associated factors (depression, schizophrenia, ADHD, etc.). For example, most (11 of 20) of the singleton probands were adopted owing to death or disability associated with neurological disease in their biological parents. To assess the relationship between identification of a P/LP variant and family history, we separated all probands into three types: simplex families in which there was only one affected proband and no 1^st^ to 3^rd^ degree relatives reported to be affected with any neurological condition (n=93); families in which the enrolled proband had no affected 1^st^ degree relatives but with one or more reported 2^nd^ or 3^rd^ degree relatives who were affected with a neurological condition (n=85); and multiplex families in which the proband had at least one first degree relative affected with a neurological condition (n=123) (Table S4). Thirty-eight probands with limited or no family history information were excluded from this analysis.

P/LP variants were found in 24 (20%) of the 123 multiplex families (20 out of 97 trios), in contrast with 35 (37.6%) of 93 simplex families (31 out of 80 trios), suggesting a P/LP identification rate that is twice as high for simplex, relative to multiplex, families. While larger sample sizes are needed to confirm this effect, the rate difference is significant whether or not all enrolled families (p=0.002) or only those sequenced as trios (p=0.008) are considered. Rates in families that were neither simplex nor multiplex (i.e., proband lacks an affected 1^st^ degree relative but has one or more affected 2^nd^ or 3^rd^ degree relatives) were intermediate, with 26% of all such families having a P/LP result (28% of trios). Of relevance to the trio/duo/singleton comparison described above, 11 of 13 (85%) singletons for which we had family history information had an affected 1^st^ degree relative, in contrast with 41% for duos and 39% for trios (Table S4). This enrichment for affected 1^st^ degree relatives likely contributed to the generally reduced rate of P/LP variants in singletons observed here.

Multiplex family findings include examples of both expected and unexpected inheritance patterns. For example, two affected male siblings were found to be hemizygous for a nonsense mutation in *PHF6* (Börjeson-Forssman-Lehmann syndrome MIM:301900) inherited from their unaffected mother. In another family, we found the proband to be compound heterozygous for two variants in *GRIK4*, with one allele inherited from each parent. Interestingly, both the mother and father of this proband report psychiatric illness, and extended family history of psychiatric phenotypes is notable. While these data are insufficient to conclude that they are indeed causative, it is plausible that the observed psychiatric phenotypes are at least partially attributable to the variation in *GRIK4* found in this family. We also observed independent *de novo* causal variants within two families. Affected siblings in family 00135 each harbored a returnable *de novo* variant in a different gene, including a VUS in *SPR* (Dystonia MIM:612716) and a pathogenic variant in *RIT1* (Noonan syndrome MIM:615355), while two probands (00075-C and 00078-C) who were second degree relatives to one another harbored independent pathogenic *de novo* variants, one each in *DDX3X* (X-linked ID MIM:300958) and *TCF20* (Table S2).

### Alternative mechanisms of disease

While the majority of DD/ID-associated genetic variation found here is predicted to lead to missense, frameshift, or nonsense effects (Figure S2B), a subset of probands harbor variants predicted to disrupt splicing, and in some cases, potentially alternative mechanisms of disease. As an example, we sequenced an affected 14-year-old girl (00003-C, Table S2) who presented with severe ID, seizures, speech delay, autism and stereotypic behaviors. WES revealed an SNV within the splice acceptor site of intron 2 in *MECP2* (c.27-6C>G, MIM:312750), identical to a previously observed *de novo* variant in a 5-year-old female with several features of Rett syndrome, but who lacked deceleration of head growth and exhibited typical growth development [23]. Laccone, et al. showed by qPCR that the variant produces a cryptic splice acceptor site that adds five nucleotides to the mRNA resulting in a frameshift (p.R9fs24X) [23]. It is likely that both the canonical and cryptic splice sites function, allowing for most *MECP2* transcripts to produce full-length protein, resulting in the milder Rett phenotype observed in the individual described here and the girl described by Laccone and colleagues [23].

In another affected proband (00126-C), we identified compound heterozygous variants in *ALG1* (Table S2). This proband has phenotypes consistent with ALG1-CDG (congenital disorder of glycosylation MIM:608540) including severe ID, hypotonia, growth retardation, microcephaly, and seizures, and was included as part of a comprehensive study of *ALG1*-associated phenotypes [24]. The paternally inherited missense mutation (c.773C>T (p.S258L)) has been previously reported as pathogenic [25], while the maternally inherited variant, which has not been observed before (c.1187+3A>G), is three bases downstream of an exon/intron junction (Figure 1A). We performed qPCR from patient blood RNA and found that intron 11 of *ALG1* is completely retained in both the proband and the mother (Figure 1A-D). The retention of intron 11 results in a stop-gain after adding 84 nucleotides (28 codons).

In a separate family consisting of affected maternal half siblings (00218-C and 00218-S, Table S2, Figure 1E) we found a variant in a canonical splice acceptor site (c.505-2A>G) of *MTOR* intron 4. The half siblings described here both have ID; the younger sibling has no seizures but has facial dysmorphism, speech delay, and autism, while his older sister exhibits seizures. We presume that the maternal half siblings inherited the splice variant from their mother, for whom DNA was not available, who was reported to exhibit seizures. We conducted qPCR and Sanger sequencing using blood-derived RNA from both siblings, finding transcripts that included an additional 134 nucleotides from the 3′ end of intron 4, ultimately leading to the addition of 20 amino acids before a stop-gain (Figure 1F-H, Figure S3). Because the stop-gain occurs early in protein translation, this splice variant likely leads to *MTOR* loss-of-function. Mutations in *MTOR* associate with a broad spectrum of phenotypes including epilepsy, hemimegalencephaly, and intellectual disability [26]. However, previously reported pathogenic variants in *MTOR* are all missense and suspected to result in gain-of-function [27]. Owing to this mechanistic uncertainty, we have classified this splice variant as a VUS. However, given the overlap between phenotypes observed in this family and previously reported families, we find this variant to be highly intriguing and suggestive that *MTOR* loss-of-function variation may also lead to disease. *MTOR* is highly intolerant of mutations in the general population (RVIS [28] score of 0.09%) supporting the hypothesis that loss-of-function is deleterious and likely leads to disease consequences.

### Proband-only versus trio sequencing

Our trio-based study design allows rapid identification of *de novo* variants, which are enriched among variants that are causally related to deleterious, pediatric phenotypes [29]. However, we also assessed to what extent our P/LP rate would differ if we had only enrolled probands. Thus, and to avoid the confounding of family history differences among trios, duos, and singletons (see above), we subjected variants found by WGS within all trio-based probands to various filtering scenarios blinded to parental status and assessed the CADD score [21] ranks of *de novo* variants previously classified as P/LP (Figure 2; Table S3). While parentally informed filters were the most sensitive and efficient (e.g., >60% of P/LP variants were the top-ranked variant among the list of all *de novo* events in each respective proband), filters defined without parental information were also effective. For example, among all rare, protein-altering (i.e., missense, nonsense, frameshift, or canonical splice-site) mutations found in genes associated with Mendelian disease via OMIM [13] or associated with DD/ID via DECIPHER [16], 20% of P/LP variants were the top-ranked variant in the given proband, most ranked among the top 5, and >80% ranked among the top 25. These data suggest that most P/LP variants could be found within probands analyzed without parental information, although additional curation time, likely in proportion to the drops in P/LP variant rank within any given filtered subset, would be required (Table S3).

In contrast to P/LP variants, VUSs would have been more difficult to identify without parental sequencing (Figure S4), owing to the fact that many VUSs do not affect genes known to associate with disease. Also, those VUSs that do affect genes known to associate with disease tended to have lesser computationally estimated effects, and therefore lower CADD ranks [21]; if they were more overtly deleterious, they would likely have been classified as P/LP. Discovery of candidate or novel disease associations, many of which are likely to eventually be shown as robust, is thus substantially more effective within trios.

### Secondary findings in participating parents

We found genetic variation unrelated to DD/ID, i.e., secondary findings, in 8.7% of parents (Table S5). One and a half percent of parents were found to harbor a P/LP variant related to a self-reported secondary condition, such as variants in *SLC22A5* that underlie a primary carnitine deficiency (MIM:212140). We also examined 56 genes identified by the ACMG as potentially harboring actionable secondary findings [14], revealing P/LP variants in 12 parents (2.0%), a rate similar to that observed in other cohorts [14, 30]. Finally, we performed a limited carrier screening assessment, identifying 28 (4.6%) parents as carriers of P/LP variation in *HBB* (Sickle cell anemia MIM:603903), *HEXA* (Tay-Sachs disease MIM:272800), or *CFTR* (Cystic fibrosis MIM:219700). We also assessed parents as mate pairs and searched for genes in which both are heterozygous for a P/LP recessive allele. These analyses yielded one parental pair (among 285 total) as carriers for variants in *ATP7B*, associated with Wilson disease (MIM:277900).

### Reanalysis of WES and WGS data

To exploit steady increases in human genetic knowledge, we performed systematic reanalyses of WES/WGS data. We approached reanalysis in three ways: 1) systematic reanalysis of old data, with the goal of reassessing each dataset every 12 months after initial analysis; 2) mining of variant prompted by new DD/ID genetic publications; and 3) use of GeneMatcher [7] to aid in the interpretation of variants in genes of uncertain disease significance.

As shown in Table 3, these efforts led to an increase in pathogenicity score for 15 variants in 17 individuals. In nine cases, a new publication became available that allowed a variant that had not been previously reported, or that was previously reported as a VUS, to be reclassified as P/LP. Three additional changes were a result of discussions facilitated by GeneMatcher [7], while the remaining upgrades resulted from reductions in filter stringency (changes to read depth and batch allele frequency) or clarification of the clinical phenotype. Among all 44 VUSs thus far identified, five (11.3%) have been upgraded. The most rapid change affected a *de novo* variant in *DDX3X*, which was upgraded from VUS to pathogenic approximately one month after initial assessment, while a *de novo* disruption of *EBF3* was upgraded from VUS to pathogenic approximately 2.5 years after initial assessment. VUSs associated with DD/ID, especially when identified via parent-proband trio sequencing, thus have considerable potential for upgrade. Additionally, of the 211 families who originally received a negative result, P/LP variation was identified for 10 (4.7%) through reanalysis. These data show that regular reanalysis of both uncertain and negative results is an effective mechanism to improve diagnostic yield.

**Table 3:**
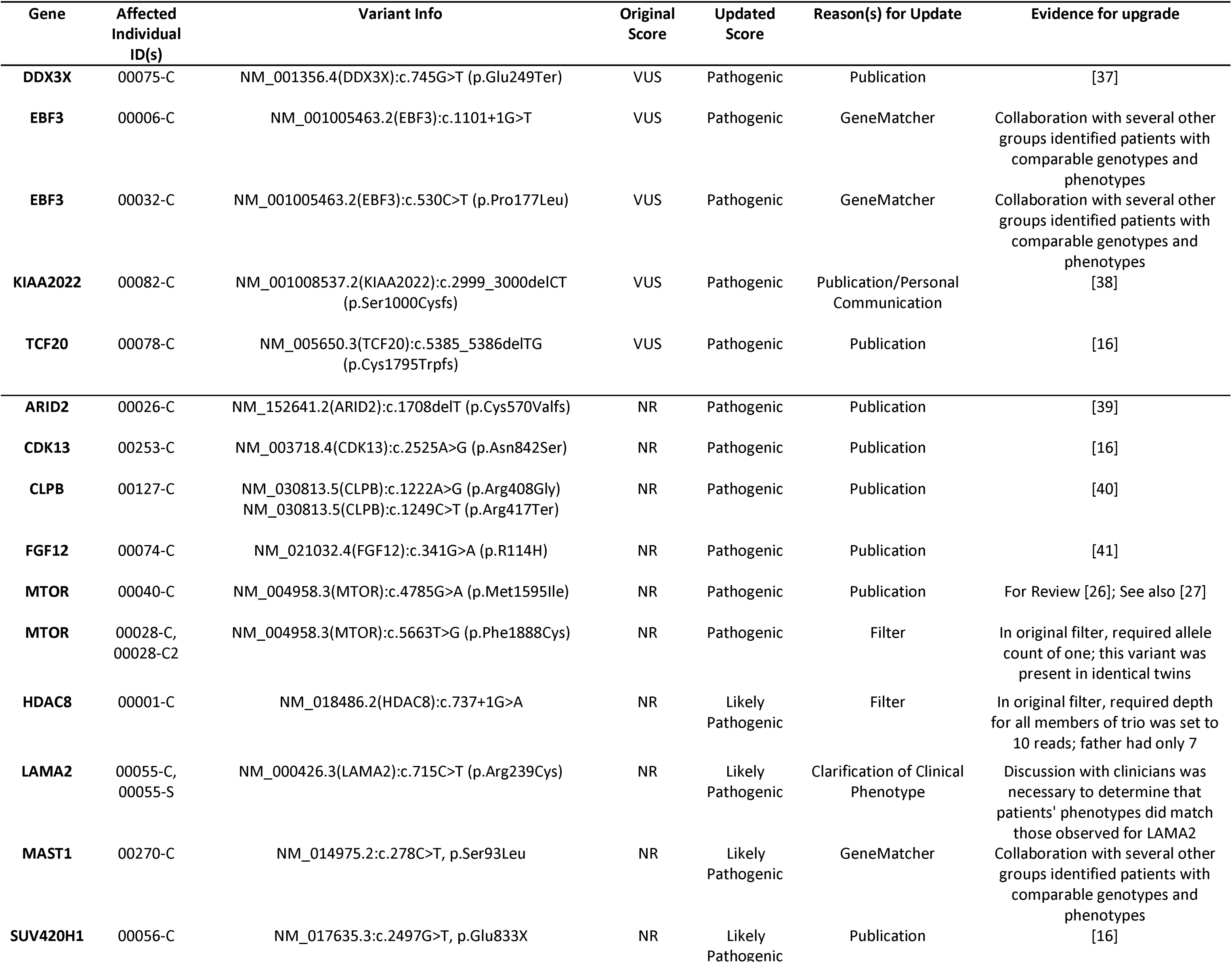
Variants with an increase in pathogenicity score due to reanalysis.

### Identification of novel candidate genes

We have identified 21 variants within 19 genes with no known disease association but which are interesting candidates. For example, in one proband (00265-C) we identified an early nonsense variant (c.2140C>T (p.R714X), CADD score 44) in *ROCK2*, with reduction of ROCK2 protein confirmed by western blot (Figure S5). ROCK2 is a conserved Rho-associated serine/threonine kinase involved in a number of cellular processes including actin cytoskeleton organization, proliferation, apoptosis, extracellular matrix remodeling and smooth muscle cell contraction, and has an RVIS [28]score placing it among the top 17.93% most intolerant genes [31]. As a second example, in two unrelated probands (00310-C and 00030-C), we identified *de novo* variation in *NBEA*, a nonsense variant at codon 2213 (of 2946, c.6637C>T (p.R2213X), CADD score 52), and a missense at codon 946 (c.2836C>T (p.H946Y), CADD score 25.6). *NBEA* is a kinase anchoring protein with roles in the recruitment of cAMP dependent protein kinase A to endomembranes near the trans-Golgi network [32]. The RVIS score [28] of *NBEA* is 0.75%.

While these variants remain VUSs, the fact that they are *de novo*, predicted to be deleterious, and affect genes under strong selective conservation in human populations, suggests they have a good chance to be disease-associated.

## DISCUSSION

We have sequenced 371 individuals with various DD/ID-related phenotypes. 27% percent of these individuals harbored a P/LP variant, most of which were *de novo* and protein-altering. We found that the P/LP yield is impacted by presence of neurological disease in family members, as our success rate drops from 38% for probands without any affected relatives to 19.5% for probands with one or more affected 1^st^ degree relatives. These data are consistent with the observation of higher causal variant yields in simplex families relative to multiplex families affected with autism [33]. It in part reflects the eased interpretation of *de novo* causal variation relative to inherited, and likely in many cases variably expressive or incompletely penetrant, causal variation (e.g., 16p12) [34].

127 probands were subject to WES and 244 were subject to WGS. The P/LP identification rate was not significantly different between the two assays when considering only SNVs or small indels (p=0.30). However, WGS is a better assay for detection of CNVs [35] and, while our patient population is depleted for large causal CNVs owing to prior array or karyotype testing, we have identified CNVs that we classified as P/LP in eight individuals.

We have also demonstrated the value of systematic reanalysis, which has thus far yielded P/LP variants for an additional 17 individuals (17% of total P/LP variants, 4.6% of total probands). Given the rates of progress in Mendelian disease genetics [36] and the development of new genomic annotations, we believe that systematic reanalysis of genomic data should become standard practice. While the costs and logistical demands for implementation at large scales are unclear, re-analysis has the potential to considerably increase diagnostic yields over time (e.g., in our study, ~8% for cases > 1 year removed from initial analysis). Furthermore, as more pathogenic coding and non-coding variants are found, the reanalysis benefit potential is largest for WGS relative to WES; the former typically has slightly better coverage of coding exons in both our data (Table S6) and previous studies [35], and re-analysis of pathogenic non-coding variation is impossible with WES.

Our data clearly suggests trio-based sequencing as more sensitive and analytically efficient than proband-only sequencing, supporting the value of trios in clinical diagnostics; as sequencing costs continue to drop, testing parents should eventually be offered routinely. Further, VUSs and novel candidates are more difficult to identify without parental sequence data, and proband-only approaches will ultimately confer less benefit in terms of discovery of new disease associations. However, current sequencing costs, when coupled to overall priorities (e.g., per-patient yield vs. total number of diagnoses) may lead to variability in decision-making about how to best allocate resources. For example, tripling per-patient sequencing costs will, under many realistic cost scenarios, lead to fewer total diagnoses within a given total budget even though the per-patient diagnostic yield is higher and curation time reduced for trios relative to singletons. Our retrospective analyses, in which we evaluated ranks of pathogenic variants under various filtering parameters, may provide useful information in making these decisions. Trade-offs in curation time, which will correlate with P/LP variant ranks, and sensitivity can be estimated empirically, in relative terms, using these data (Figure 2; Table S3).

Variation detected through our studies has already helped lead to the discovery of at least one new disease association, as we identified two patients that harbor *de novo* variants in *EBF3*, a highly conserved transcription factor involved in neurodevelopment that is relatively intolerant to mutations in the general population (RVIS [28]: 6.78%). Through collaboration with other researchers via GeneMatcher [7], we were able to identify a total of 10 DD/ID-affected individuals who harbor *EBF3* variants, supporting that *de novo* disruption of *EBF3* function leads to neurodevelopmental phenotypes [37]. It is our hope that the other VUSs described here, shared via ClinVar [6] and GeneMatcher [7], will also help to facilitate new associations.

## CONCLUSIONS

We have demonstrated the benefits of genomic sequencing to identify disease-associated variation in probands with developmental disabilities who are otherwise lacking a precise clinical diagnosis. Indeed, by combining genomic breadth with resolution capable of detecting SNVs, indels, and CNVs in a single assay, WGS is a highly effective choice as the first diagnostic test, rather than last resort, for unexplained developmental disabilities. The ability for WGS to serve as a single-assay replacement for WES and microarrays underscores its value as a frontline test. Furthermore, the benefits and effectiveness of WGS testing is likely to grow over time both by accelerating research (for example into the discovery of smaller pathogenic CNVs and pathogenic SNVs outside of coding exons), and by facilitating more effective reanalysis, a process which we show to be an essential component to maximize diagnostic yield.

## ABBREVIATIONS

WGS, whole genome sequencing; WES, whole exome sequencing; CSER, Clinical Sequencing Exploratory Research; CNV, copy number variant; DD/ID, Developmental delay/Intellectual disability; P/LP, Pathogenic/likely pathogenic; VUS, variant of uncertain significance

## DECLARATIONS

*Ethics approval and consent to participate:* Review boards at Western (20130675) and the University of Alabama at Birmingham (X130201001) approved and monitored this study. *Consent for publication:* A parent or legal guardian was required to give consent to participate in the study and inclusion of their data for publication, and assent was obtained from those children who were capable. *Availability of data and material:* The genomic data generated for this work is available through dbGAP [5], ClinVar [6], and GeneMatcher [7]. *Competing interests:* The authors declare that they have no competing interests. *Funding:* This work was supported by grants from the US National Human Genome Research Institute (NHGRI; UM1HG007301) and the National Cancer Institute (NCI; R01CA197139). *Author contributions:* GMC, RMM, GSB, NEL and KBB designed the study and guided its implementation. KMB, MDA, CRF, SMH, and MLT performed genomic analyses to identify disease-linked variation, prepared variant reports, and oversaw variant review. DEG called CNVs from WGS data. DEG, BTW and JSW contributed to bioinformatics analyses. ASN contributed to data acquisition. WVK, KME, SS, EJL, and EMB recruited study participants, collected blood samples and clinical information, and returned results. KBB, CAR, GSB aided in variant review. KLE and JNC performed functional validation studies. KMB, MDA, CRF, SMH, MLT, and GMC wrote the manuscript. All authors contributed to and approved the manuscript.

## Acknowledgements

We are grateful to the patients and their families who contributed to this study. We thank the HudsonAlpha Software Development and Informatics team and the Genome Sequencing Center who contributed to data acquisition and analysis. We would also like to thank Dr. Jeremy Herskowitz in the Department of Neurology at University of Alabama at Birmingham for discussions about ROCK2.

## REFERENCES

1. Boyle CA, Boulet S, Schieve LA, Cohen RA, Blumberg SJ, Yeargin-Allsopp M, et al. Trends in the prevalence of developmental disabilities in US children, 1997-2008. Pediatrics. 2011;127:1034–42.

2. Sun F, Oristaglio J, Levy SE, Hakonarson H, Sullivan N, Fontanarosa J, et al. Genetic Testing for Developmental Disabilities, Intellectual Disability, and Autism Spectrum Disorder. Rockville (MD): Agency for Healthcare Research and Quality (US). 2015 Jun (Technical Briefs, No. 23) Available from: https://www.ncbi.nlm.nih.gov/books/NBK304462/.

3. Stavropoulos D, Merico D, Jobling R, Bowdin S. Whole-genome sequencing expands diagnostic utility and improves clinical management in paediatric medicine. npj Genomic Medicine. 2016;1:1–9.

4. Green RC, Goddard KA, Jarvik GP, Amendola LM, Appelbaum PS, Berg JS, et al. Clinical Sequencing Exploratory Research Consortium: Accelerating Evidence-Based Practice of Genomic Medicine. Am J Hum Genet. 2016;98:1051–66.

5. Mailman MD, Feolo M, Jin Y, Kimura M, Tryka K, Bagoutdinov R, et al. The NCBI dbGaP database of genotypes and phenotypes. Nat Genet. 2007;39:1181–6.

6. Landrum MJ, Lee JM, Benson M, Brown G, Chao C, Chitipiralla S, et al. ClinVar: public archive of interpretations of clinically relevant variants. Nucleic Acids Res. 2016;44:D862–8.

7. Sobreira N, Schiettecatte F, Valle D, Hamosh A. GeneMatcher: a matching tool for connecting investigators with an interest in the same gene. Hum Mutat. 2015;36:928–30.

8. Li H, Durbin R. Fast and accurate short read alignment with Burrows-Wheeler transform. Bioinformatics. 2009;25:1754–60.

9. DePristo MA, Banks E, Poplin R, Garimella KV, Maguire JR, Hartl C, et al. A framework for variation discovery and genotyping using next-generation DNA sequencing data. Nat Genet. 2011;43:491–8.

10. Manichaikul A, Mychaleckyj JC, Rich SS, Daly K, Sale M, Chen WM. Robust relationship inference in genome-wide association studies. Bioinformatics. 2010;26:2867–73.

11. Zhu M, Need AC, Han Y, Ge D, Maia JM, Zhu Q, et al. Using ERDS to infer copy-number variants in high-coverage genomes. Am J Hum Genet. 2012;91:408–21.

12. Miller CA, Hampton O, Coarfa C, and Milosavljevic A. ReadDepth: a parallel R package for detecting copy number alterations from short sequencing reads. PLoS One. 2011;6:e16327.

13. Hamosh A, Scott AF, Amberger JS, Bocchini CA, and McKusick VA. Online Mendelian Inheritance in Man (OMIM), a knowledgebase of human genes and genetic disorders. Nucleic Acids Res. 2005;33:D514–7.

14. Green RC, Berg JS, Grody WW, Kalia SS, Korf BR, Martin CL, et al. ACMG recommendations for reporting of incidental findings in clinical exome and genome sequencing. Genet Med. 2013;15:565–74.

15. Lek M, Karczewski KJ, Minikel EV, Samocha KE, Banks E, Fennell T, et al. Analysis of protein-coding genetic variation in 60,706 humans. Nature. 2016;536:285–91.

16. Deciphering Developmental Disorders Study. Prevalence and architecture of de novo mutations in developmental disorders. Nature. 2017; doi:10.1038/nature21062.

17. Epi4K Consirum, Epilepsy Phenome/Genome Project, Allen AS, Berkovic SF, Cossette P, Delanty N, et al. De novo mutations in epileptic encephalopathies. Nature. 2013;501:217–21.

18. Karaca E, Harel T, Pehlivan D, Jhangiani SN, Gambin T, Coban Akdemir Z, et al. Genes that Affect Brain Structure and Function Identified by Rare Variant Analyses of Mendelian Neurologic Disease. Neuron. 2015;88:499–513.

19. Samocha KE, Robinson EB, Sanders SJ, Stevens C, Sabo A, McGrath LM, et al. A framework for the interpretation of de novo mutation in human disease. Nat Genet. 2014;46:944–50.

20. Richards S, Aziz N, Bale S, Bick D, Das S, Gastier-Foster J, et al. Standards and guidelines for the interpretation of sequence variants: a joint consensus recommendation of the American College of Medical Genetics and Genomics and the Association for Molecular Pathology. Genet Med. 2015;17:405–24.

21. Kircher M, Witten DM, Jain P, O’Roak BJ, Cooper GM, Shendure J. A general framework for estimating the relative pathogenicity of human genetic variants. Nat Genet. 2014;46:310–5.

22. Kohler S, Doelken SC, Mungall CJ, Bauer S, Firth HV, Bailleul-Forestier I, et al. The Human Phenotype Ontology project: linking molecular biology and disease through phenotype data. Nucleic Acids Res. 2014;42:D966–74.

23. Laccone F, Huppke P, Hanefeld F, Meins M. Mutation spectrum in patients with Rett syndrome in the German population: Evidence of hot spot regions. Hum Mutat. 2001;17:183–90.

24. Ng BG, Shiryaev SA, Rymen D, Eklund EA, Raymond K, Kircher M, et al. ALG1-CDG:Clinical and Molecular Characterization of 39 Unreported Patients. Hum Mutat. 2016;37:653–60.

25. Dupre T, Vuillaumier-Barrot S, Chantret I, Sadou Yaye H, Le Bizec C, Afenjar A, et al. Guanosine diphosphate-mannose:GlcNAc2-PP-dolichol mannosyltransferase deficiency (congenital disorders of glycosylation type Ik): five new patients and seven novel mutations. J Med Genet. 2010;47:729–35.

26. Baulac S. mTOR signaling pathway genes in focal epilepsies. Prog Brain Res. 2016;226:61–79.

27. Moller R, Wechkuysen S, Chipaux M, Marsan E, Taly V, Bebin ME, et al. Germline and somatic mutations in MTOR gene in focal cortical dysplasia and epilepsy. Neurology: Genetics. In press;2:e118.

28. Petrovski S, Wang Q, Heinzen EL, Allen AS, Goldstein DB. Genic intolerance to functional variation and the interpretation of personal genomes. PLoS Genet. 2013;9:e1003709.

29. Vissers LE, de Ligt J, Gilissen C, Janssen I, Steehouwer M, de Vries P, et al. A de novo paradigm for mental retardation. Nat Genet. 2010;42:1109–12.

30. Johnston JJ, Rubinstein WS, Facio FM, Ng D, Singh LN, Teer JK, et al. Secondary variants in individuals undergoing exome sequencing: screening of 572 individuals identifies high-penetrance mutations in cancer-susceptibility genes. Am J Hum Genet. 2012;91:97–108.

31. Loirand G. Rho Kinases in Health and Disease: From Basic Science to Translational Research. Pharmacol Rev. 2015;67:1074–95.

32. Castermans D, Wilquet V, Parthoens E, Huysmans C, Steyaert J, Swinnen L, et al. The neurobeachin gene is disrupted by a translocation in a patient with idiopathic autism. J Med Genet. 2003;40:352–6.

33. Klei L, Sanders SJ, Murtha MT, Hus V, Lowe JK, Willsey AJ, et al. Common genetic variants, acting additively, are a major source of risk for autism. Mol Autism. 2012;3:9.

34. Quintans B, Ordonez-Ugalde A, Cacheiro P, Carracedo A, Sobrido MJ. Medical genomics: The intricate path from genetic variant identification to clinical interpretation. Appl Transl Genom. 2014;3:60–7.

35. Belkadi A, Bolze A, Itan Y, Cobat A, Vincent QB, Antipenko A, et al. Whole-genome sequencing is more powerful than whole-exome sequencing for detecting exome variants. Proc Natl Acad Sci U S A. 2015;112:5473–8.

36. Chong JX, Buckingham KJ, Jhangiani SN, Boehm C, Sobreira N, Smith JD, et al. The Genetic Basis of Mendelian Phenotypes: Discoveries, Challenges, and Opportunities. Am J Hum Genet. 2015;97:199–215.

37. Harms FL, K.M. G, Hardigan AA, Kortum F, Shukla A, Alawi M, et al. Mutations in EBF3 disturb transcriptional profiles and underlie a novel syndrome of intellectual disability, ataxia and facial dysmorphism. BioRxiv. 2016; https://doi.org/10.1101/067454.

38. Deciphering Developmental Disorders Study. Large-scale discovery of novel genetic causes of developmental disorders. Nature. 2015;519:223–8.

39. Farach LS, Northrup H. KIAA2022 nonsense mutation in a symptomatic female. Am J Med Genet A. 2016;170:703–6.

40. Shang L, Cho MT, Retterer K, Folk L, Humberson J, Rohena L, et al. Mutations in ARID2 are associated with intellectual disabilities. Neurogenetics. 2015;16:307–14.

41. Wortmann SB, Zietkiewicz S, Kousi M, Szklarczyk R, Haack TB, Gersting SW, et al. CLPB mutations cause 3-methylglutaconic aciduria, progressive brain atrophy, intellectual disability, congenital neutropenia, cataracts, movement disorder. Am J Hum Genet. 2015;96:245–57.

42. Siekierska A, Isrie M, Liu Y, Scheldeman C, Vanthillo N, Lagae L, et al. Gain-of-function FHF1 mutation causes early-onset epileptic encephalopathy with cerebellar atrophy. Neurology. 2016;86:2162–70.

43. O’Leary NA, Wright MW, Brister JR, Ciufo S, Haddad D, McVeigh R, et al. Reference sequence (RefSeq) database at NCBI: current status, taxonomic expansion, and functional annotation. Nucleic Acids Res. 2016;44:D733–45.

44. Genomes Project C, Auton A, Brooks LD, Durbin RM, Garrison EP, Kang HM, et al. A global reference for human genetic variation. Nature. 2015;526:68–74.

45. Exome Variant Server, NHLBI GO Exome Sequencing Project (ESP). [cited 2013; Available from: http://evs.gs.washington.edu/EVS.

